# Detecting Local Genetic Correlations with Scan Statistics

**DOI:** 10.1101/808519

**Authors:** Hanmin Guo, James J. Li, Qiongshi Lu, Lin Hou

## Abstract

Genetic correlation analysis has quickly gained popularity in the past few years and provided insights into the genetic etiology of numerous complex diseases. However, existing approaches oversimplify the shared genetic architecture between different phenotypes and cannot effectively identify precise genetic regions contributing to the genetic correlation. In this work, we introduce LOGODetect, a powerful and efficient statistical method to identify small genome segments harboring local genetic correlation signals. LOGODetect automatically identifies genetic regions showing consistent associations with multiple phenotypes through a scan statistic approach. It uses summary association statistics from genome-wide association studies (GWAS) as input and is robust to sample overlap between studies. Applied to five phenotypically distinct but genetically correlated psychiatric disorders, we identified 49 non-overlapping genome regions associated with multiple disorders, including multiple hub regions showing concordant effects on more than two disorders. Our method addresses critical limitations in existing analytic strategies and may have wide applications in post-GWAS analysis.

## Introduction

Genome-wide association studies (GWASs) have been carried out for numerous complex traits and diseases, identifying tens of thousands of single-nucleotide polymorphisms (SNPs) associated with these phenotypes. However, our understanding of most traits’ genetic basis remains incomplete, in part due to the limited power and interpretability of the traditional GWAS approach that correlates one trait with one SNP at a time. Recently, statistical methods that jointly model multiple phenotypes have quickly gained popularity in human genetics research^1–3^. Leveraging pervasive pleiotropy in the human genome, these methods enhanced the statistical power to identify genetic associations^1, 4–7^, improved the accuracy of genetic risk prediction^8,9^, revealed novel genetic sharing across diverse phenotypes^10–12^, and provided great insights into the genetic basis of a variety of diseases and traits^13,14^.

Genetic similarity between traits can be modeled at different scales. Methods that identify SNPs associated with multiple phenotypes have achieved some success^15–17^. However, most complex human traits are highly polygenic, with top SNPs showing weak to moderate effects^18,19^. Thus, single SNP-based methods may not be sufficient to characterize the full landscape of genetic similarity. An alternative approach is to estimate the genetic correlation between different traits^10,12,20,21^. These methods effectively utilize genome-wide genetic data, including SNPs that do not reach statistical significance in GWAS, to quantify the overall genetic sharing between two traits. In addition, recent methodological advances have enabled estimation of genetic correlation with GWAS summary statistics^10,11,22^, making these approaches widely applicable to a large number of complex phenotypes. With these advances, genetic correlation analysis has become a routine procedure in post-GWAS analysis and was implemented in almost all large-scale GWASs published in the past few years.

However, despite improved statistical power and wide applications, genetic correlation approaches fail to provide detailed, mechanistic insights due to its oversimplification of complex genetic sharing into a single metric. Two recent methods improved genetic correlation analysis by providing local^12^ and annotation-stratified estimates^11^. However, these methods rely on strong prior evidence about which local region or functional annotation to investigate. When applied to hypothesis-free scans, statistical power is substantially reduced. In this work, we introduce LOGODetect (LOcal Genetic cOrrelation Detector), a novel method that uses scan statistics to identify genome segments harboring local genetic correlation between two complex traits. Compared to other methods, LOGODetect does not pre-specify candidate regions of interest, and instead, automatically detects regions with shared genetic components with great resolution and statistical power. In addition, LOGODetect only uses GWAS summary statistics as input and is robust to sample overlap between GWASs. We demonstrate its performance through extensive simulations and analysis of well-powered GWASs for five distinct but genetically correlated psychiatric disorders^23,24^. Our analysis implicates a collection of hub regions in the genome that underlie the risk for several of these disorders.

## Results

### Method overview

Our goal is to identify genome segments showing consistent association patterns with two different traits. Here, we provide an overview of our approach and the technical details are discussed in **Methods**. We propose the following scan statistic

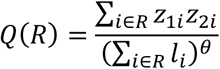

to quantify the extent of local genetic similarity in a genome region, where *R* is the index set for all SNPs in the region, *z*_1*i*_ and *z*_2*i*_ are the association z-scores for the *i*-th SNP with two traits, *l*_*i*_ is the linkage disequilibrium (LD) score for the *i*-th SNP,^10^ and *θ* controls the impact of LD. *Q*(*R*) is a LD score-weighted inner product of local z-scores from two GWASs and is conceptually similar to local genetic correlation - regions with high absolute values of *Q*(*R*) show concordant association patterns across multiple SNPs in the region and the sign of *Q*(*R*) shows if the correlation is positive or negative. We search for genome segments with the highest |*Q*(*R*)| values by scanning the genome while allowing the segment size to vary (**Figure 1**). Statistical evidence of genetic sharing is assessed using a Monte Carlo approach.

**Figure 1.**
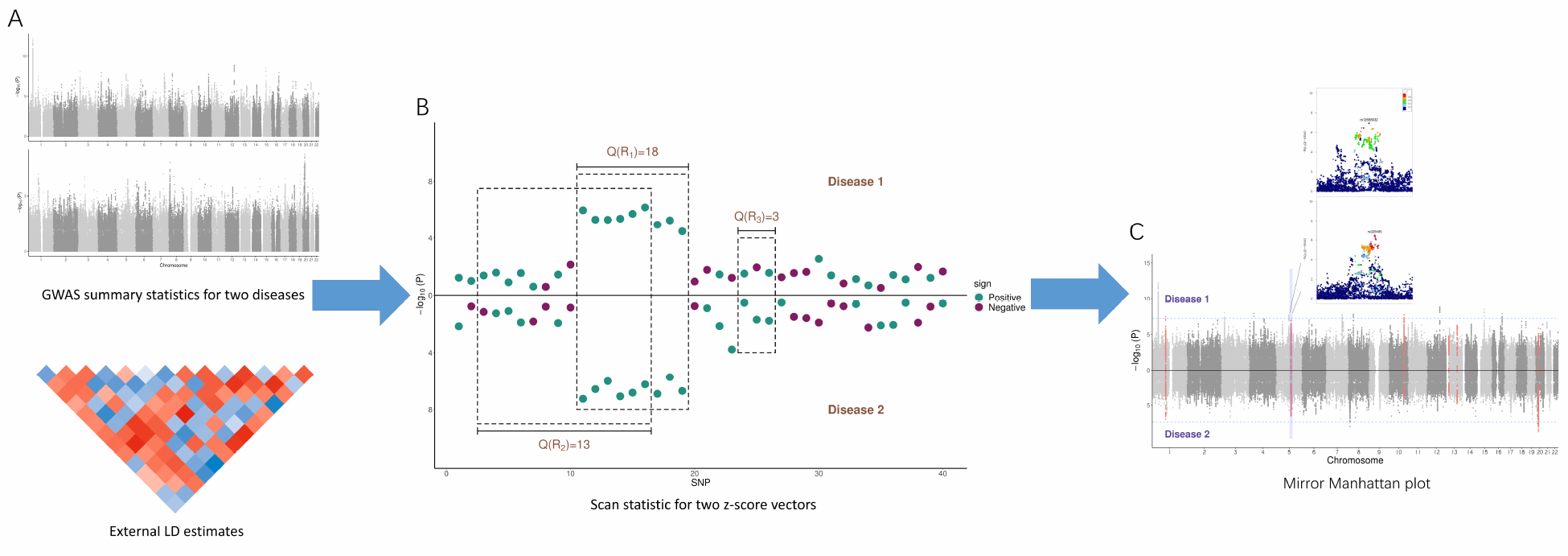
LOGODetect workflow. (A) The inputs of LOGODetect include GWAS summary statistics for two traits and a reference panel for LD estimation. (B) Scan statistic is defined over a region, as the LD-weighted inner product of two z-score vectors in this region. A large absolute value of the scan statistic would hint at local genetic correlation. (C) LOGODetect identifies genome segments showing consistent associations with two different traits.

### Simulation results

We conducted simulations to assess the type I error of our approach using 15,918 samples from the Wellcome Trust Case Control Consortium (WTCCC). First, we simulated phenotypes under an infinitesimal model in which genetic effects were assumed to be the same for all SNPs. In addition, we also investigated a more realistic genetic architecture with different levels of genetic effects - we attributed 30% of the trait heritability to 3% of randomly selected SNPs, while the remaining SNPs explain 70% of the total heritability. The family-wise type I error rate of our method was well-controlled as the heritability of each trait ranged between 0.01 to 0.9 (**Supplementary Table 1)** and under strong heritability enrichment (**Supplementary Table 2**), suggesting that LOGODetect is statistically robust to diverse genetic architecture. We also assessed the statistical power of LOGODetect under various settings (**Figure 2** and **Supplementary Figures 1-3**). Three different metrics were used to quantify the statistical power (see **Methods**). With smaller values of θ, LOGODetect tends to find long segments, achieving greater point detection rate (**Figure 2A**). On the contrary, LOGODetect identified many short segments with decreased point detection rate when greater values of θ was used. LOGODetect obtains a larger G-score with smaller *θ* (**Methods**), and in particular, the G-score is almost the same for *θ* = 0.3, 0.4 or 0.5. Overall, LOGODetect worked reasonably well in all three measures when *θ* was set to 0.5. As a result, we recommend to use 0.5 as the choice of *θ* in practice.

**Figure 2.**
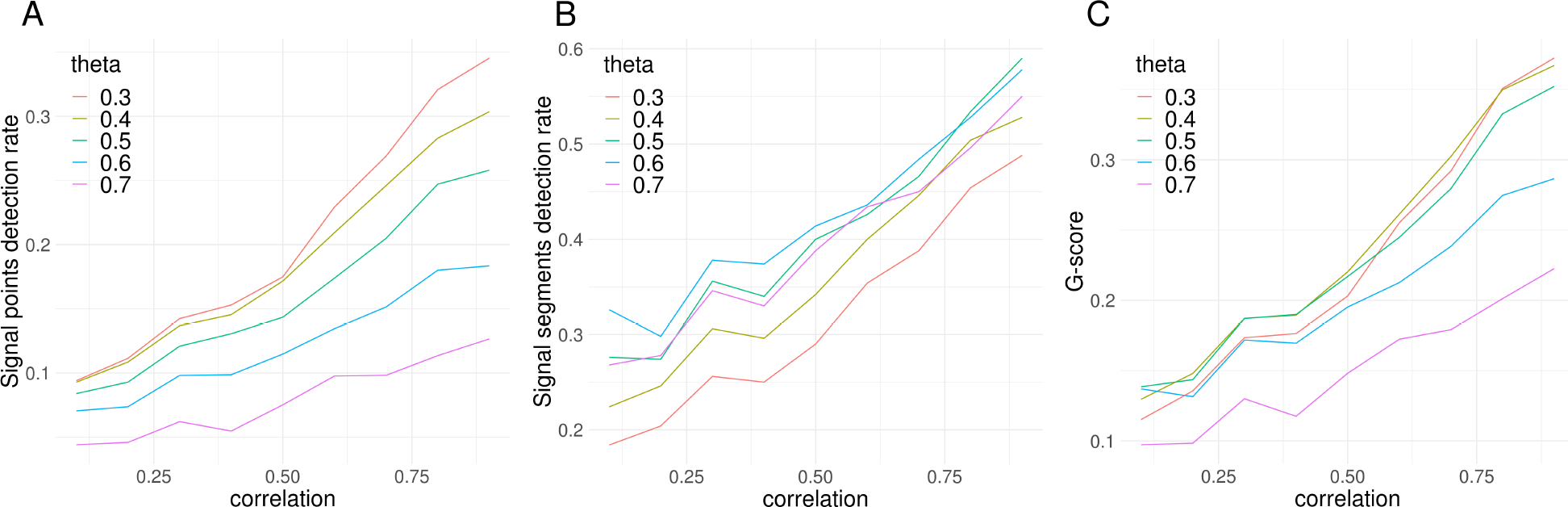
Assessment of statistical power. The Y-axis shows statistical power assessed by three different measures: (A) signal points detection rate, (B) signal segments detection rate, and (C) G-score. The X-axis shows the correlation of SNPs’ effects on two traits in the genome region.

### Application to five psychiatric disorders

Previous studies have revealed pervasive pleiotropy^25–27^ and genetic covariance^28–31^ among psychiatric disorders. However, there is limited understanding of the specific genetic loci contributing to multiple disorders. We applied LOGODetect to study the pairwise local genetic correlation between five psychiatric disorders (**Supplementary Table 3**): bipolar disorder (BIP; n=51,710), schizophrenia (SCZ; n=105,318), major depressive disorder (MDD; n=173,005), attention-deficit/hyperactivity disorder (ADHD; n=53,293), and autism spectrum disorder (ASD; n=46,350), using summary statistics from the latest GWASs.^32–36^ In total, we identified 66 regions (49 non-overlapping segments) showing concordant associations with multiple disorders (FDR < 0.05; **Figure 3** and **Supplementary Figures 4-9**). 65 of the 66 regions showed positive correlations. Size of the identified genome segments varied from 24 KB to 1.6 MB (**Supplementary Figure 10**). The number of significant segments identified in our analysis is proportional to the genetic correlation between each pair of disorders (**Supplementary Figure 11**; correlation r=0.68). We identified 33 shared genome segments for BIP and SCZ (**Figure 3B**; genetic correlation r_g_=0.68, p=9.14e−87), 4 shared regions for BIP and MDD (**Supplementary Figure 4**; r_g_ =0.42, p=4.33e−17), and 11 regions for SCZ and MDD (**Supplementary Figure 5**; r_g_=0.40, p=7.36e−33), which is consistent with the strong genetic overlap between these disorders^28, 37–39^. Additionally, studies have suggested correlated familial genetic liabilities among MDD, ADHD, and ASD.^29,30,40^ LOGODetect identified 5 regions shared by MDD and ADHD (**Supplementary Figure 7**; r_g_ =0.61, p=1.72e−15), 4 regions for MDD and ASD (**Supplementary Figure 8**; r_g_ =0.55, p=1.41e−15), and 6 regions for ADHD and ASD (**Supplementary Fig 9**; r_g_=0.42, p=4.46e−10). Overall, we identified strong genetic sharing (higher genetic correlation and more shared genome segments) among SCZ, BIP, and MDD and among MDD, ASD, and ADHD. Sharing between these two clusters was relatively weaker.

**Figure 3.**
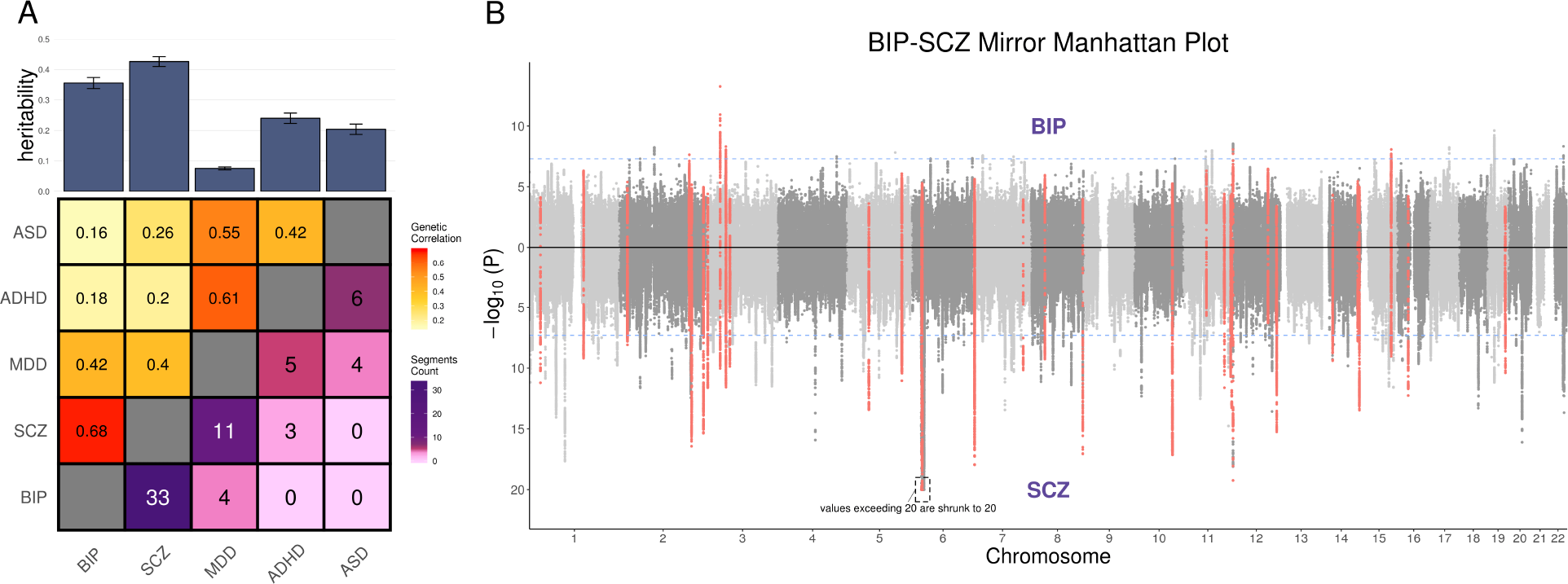
LOGODetect identifies genome regions contributing to multiple psychiatric disorders. (A) Heatmap shows the genetic correlation estimates (upper triangle) and the number of segments with local genetic correlation identified by LOGODetect (lower triangle) between five psychiatric disorders; Barplot shows the heritability estimates and standard errors of five disorders. (B) Mirrored Manhattan plot for BIP and SCZ. The 33 shared genome regions identified by LOGODetect are highlighted in red. One locus on chromosome 6 with − log_10_ *P* > 20 in SCZ is truncated at 20 for visualization purpose.

### Tissue enrichment of hub regions shared by psychiatric disorders

We used GenoSkyline-Plus tissue-specific functional annotations^41^ to investigate the functional relevance of the genomic regions found to harbor local genetic correlations among five psychiatric disorders. First, we tested the five disorders’ heritability enrichment in the predicted functional genome of 66 tissue and cell types (**Supplementary Table 6**; **Methods**).^10^ Eight tissue and cell types not enriched (p>0.01) for the heritability of any disorder were removed from the analysis. We used permutation tests to assess the enrichment of genome regions shared by multiple disorders in the remaining 58 annotation tracks. Genome regions identified by LOGODetect were significantly enriched in multiple brain regions including anterior caudate (enrichment=1.82, p=3.00e−4), cingulate gyrus (enrichment=1.94, p=2.00e−4), inferior temporal lobe (enrichment=2.00, p=3.00e−4), angular gyrus (enrichment=1.96, p=3.00e−4), and dorsolateral prefrontal cortex (enrichment=2.01, p=3.00e−4) (**Figure 4**). In addition to brain tissues, regions shared by psychiatric disorders were also enriched in pancreatic islets (enrichment=2.15, p=2.00e−4) and mononuclear cells from peripheral blood (enrichment=2.08, p=3.00e−4). Of note, annotated functional regions in these tissues have substantial overlaps with annotations of brain tissues (**Figure 4B**). After conditioning on functional regions in the brain, the enrichment in pancreatic islets was substantially reduced (enrichment=1.03, p=0.40; **Figure 4C**), while enrichment in mononuclear cells remained significant (enrichment=1.67, p=0.02).

**Figure 4.**
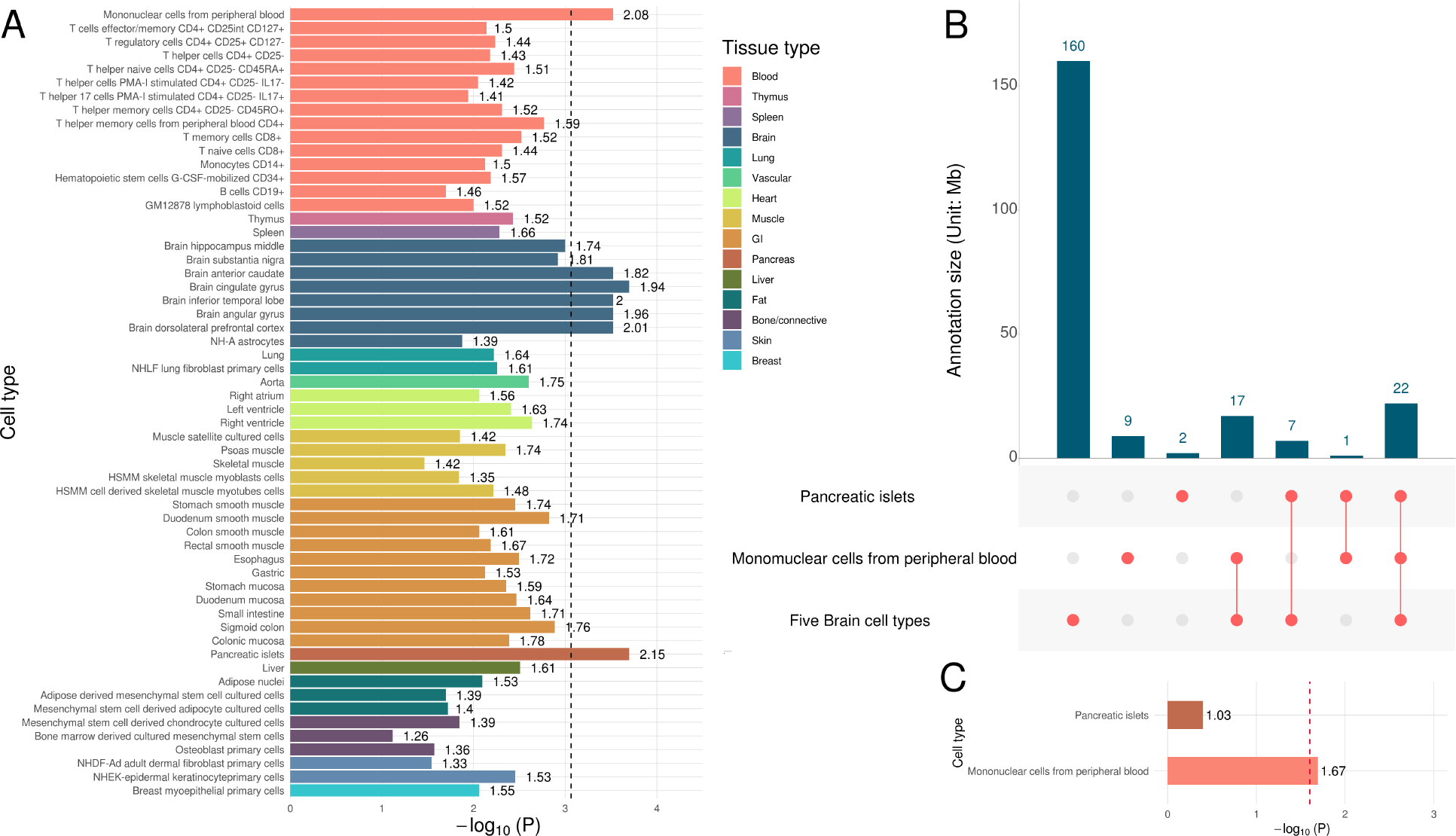
Tissue-specific enrichment of genome regions conferring risk for multiple psychiatric disorders. (A) Permutation test results over 58 cell-type specific annotations. Fold enrichment is labeled next to each bar. (B) The overlap of predicted functional regions in pancreatic islets, mononuclear cells from peripheral blood, and five brain regions. (C) Enrichment in the predicted functional regions in pancreatic islets and mononuclear cells from peripheral blood after conditioning on the annotation overlap with brain regions.

### Hub regions contributing to multiple psychiatric disorders

Next, we investigated hub regions shared by more than two disorders. Among the 49 non-overlapping genome regions identified in our analysis, 8 regions were identified in two different disorder pairs, 3 regions were identified in three pairs, and 1 region was identified in four pairs (**Supplementary Table 4**). The 4 regions identified in at least three pair-wise analyses are summarized in **Figure 5**. These hub regions show consistent associations with multiple psychiatric disorders and can potentially reveal key mechanisms and pathways underlying the shared genetics across disorders.

**Figure 5.**
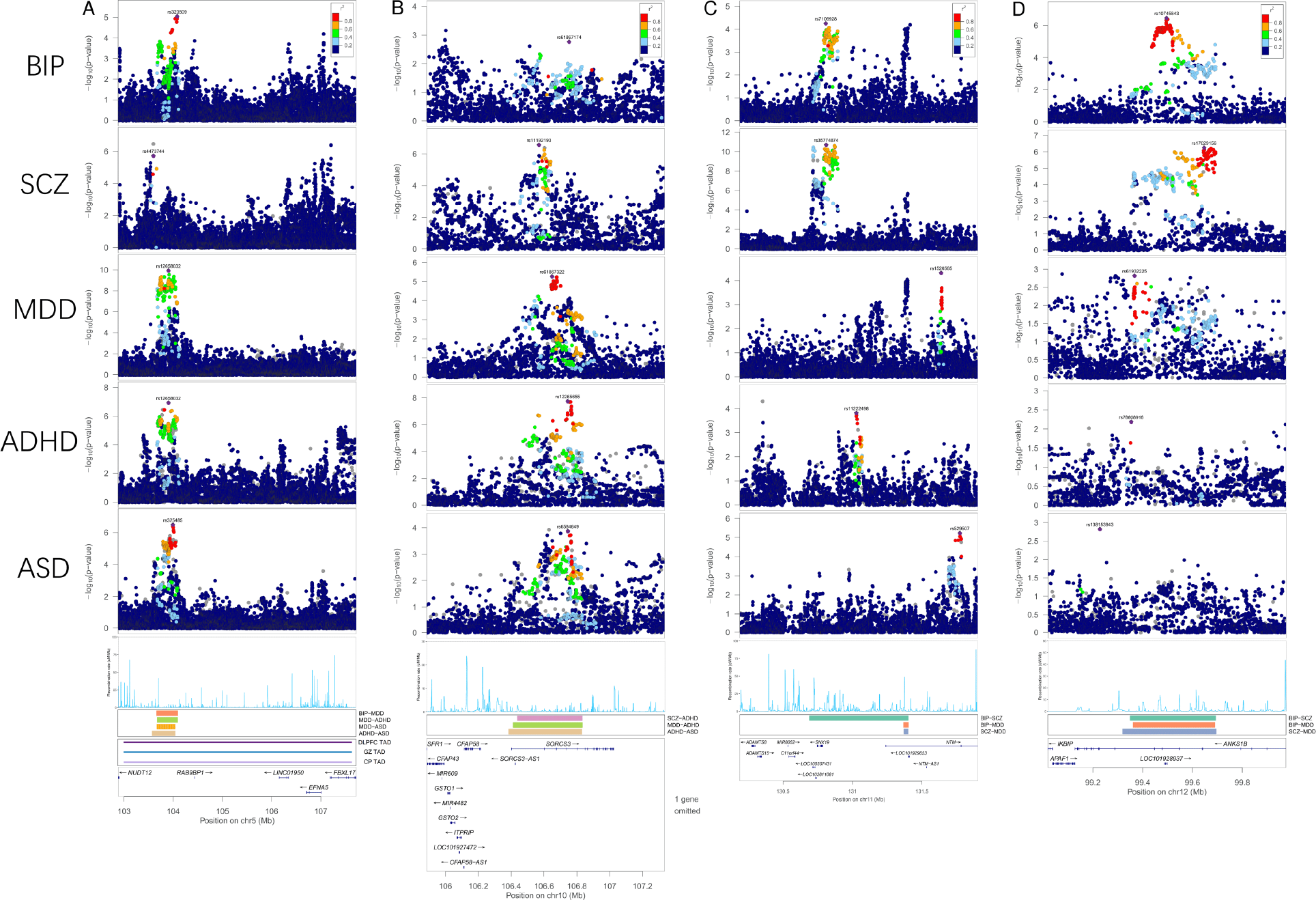
Putative target genes for four hub regions shared by more than two psychiatric disorder. (A) shows the significant region in chr5 shared by four disorder pairs, (B-D) shows there significant regions, each is shared by three disorder pairs. Locuszoom plot, recombination rate, and the gene names are provided. The colored band denote the location of the significant region and which disorder pair is detected in. TAD in dorsolateral prefrontal cortex (DLPFC) in adult brain and TAD in the germinal zone (GZ) and postmitoticzone cortical plate (CP) in the developing fetal brain are shown as long solid line in panel (A).

The region showing significant correlation between BIP-MDD (p=2.00e−4; q=0.041), MDD-ADHD (p=2.00e−4; q=0.041), MDD-ASD (p=2.00e−4; q=0.041), and ADHD-ASD (p=6.00e−4; q=0.061) is a locus spanning 500 KB on chromosome 5 (**Supplementary Table 5**). We note that this is an intergenic region but SNPs in this region have previously reached genome-wide significance in the MDD GWAS^34^ (lead SNP rs12658032; p=1.18e−10). Additionally, SNPs at this locus showed consistent associations with BIP (lead SNP rs323509; p=8.94e−6), ADHD (lead SNP rs12658032; p=1.15e−7), and ASD (lead SNP rs325485; p=3.25e−7). There are also suggestive associations with SCZ (lead SNP rs4473744; p=1.88e−6) but the lead SNP is not in LD with SNPs in the specific genome segment implicated in our analysis. Interestingly, although the closest protein-coding gene *NUDT12* is 700 KB away, this region is located in a large topological associating domain (TAD; 4.8 MB) that is conserved in adult and fetal brains (**Figure 5A**; **Methods**)^42,43^. Three genes, *EFNA5*, *NUDT12*, and *FBXL17*, are located in the TAD. We also note that the genome region identified by LOGODetect overlapped with *RP11-6N13.1*, a noncoding RNA exclusively expressed in the testis tissue. Multiple eQTLs for *RP11-6N13.1* are located in the region (lead SNP rs416223; p=1.25e−13). Although there is no direct evidence suggesting this noncoding RNA is linked to psychiatric disorders, it remains a hypothesis worth investigating in the future.

We also identified 3 additional hub regions, each shared by 3 pairs of disorders (**Supplementary Table 4**). The locus on chromosome 10 spans 450 KB and showed significant correlations between SCZ-ADHD, MDD-ADHD, and ADHD-ASD (**Figure 5B**). The genome regions identified at this locus largely overlaps with *SORCS3*, a previously implicated risk gene for MDD and ADHD.^35,44,45^ The second hub region is located on chromosome 11, spanning 715 KB. There are multiple independent association peaks in this region (**Figure 5C**) and it was significantly correlated between BIP-SCZ, BIP-MDD, and SCZ-MDD in our anlaysis. Two genes, *NTM* and *SNX19*, are located near this region. The third hub region spans 375 KB on chromosome 12 (**Figure 5D**). It showed concordant associations between BIP-SCZ, BIP-MDD, and SCZ-MDD. This region is located in *ANKS1B*, a significant gene in the genome-wide pharmacogenomic analysis of antipsychotic drug (AP) response in SCZ^46,47^.

## Discussion

Through simulations and analyses of GWAS data, we demonstrated that our method effectively identified genetic regions that may be shared across multiple complex traits with high resolution and statistical power. Applied to well-powered GWASs for five phenotypically distinct but genetically correlated psychiatric disorders, LOGODetect identified numerous shared genomic regions including hub regions that showed consistent effects for more than two disorders. Three genes (i.e. *EFNA5*, *NUDT12*, and *FBXL17*) are located in the same TAD with the hub region on chromosome 5 (**Figure 5A**). *EFNA5*, also known as *Ephrin-A5*, interacts with Eph receptors and plays a critical role in accurate guidance of cell or axon movement and synapse development in the nervous system^48–50^. It is highly expressed in various brain areas and was found to regulate the formation of the ascending midbrain dopaminergic pathways^51^ which are involved in social interactions and reward^52^. Ephrin-A5 knockout mice model suggested that *ephrin-A5* plays an important role in the normal development of central monoaminergic pathways^53^, whose alteration has been linked to SCZ, MDD, and ADHD^54,55^. In addition, *ephrin-A5* knockout mice shared some similarities in the developmental delays seen in children diagnosed with ADHD^53^. Based on the literature support and TADs derived from adult and fetal brain hi-C data, *EFNA5* is a strong candidate gene that may explain the link of this hub region with psychiatric disorders, although we do not rule out the possible involvement of *NUDT12* and *FBXL17*.

The hub regions shared by 3 pairs of disorders also overlapped with a handful of interesting candidate genes. *SORCS3* (**Figure 5B**) is highly expressed in the CA1 region of the hippocampus, and is involved in synaptic depression and spatial learning ability^56,57^. It is also known to play an important role in protein networks associated with *PICK1*, *NGF*, and *PDGF-BB*^58,59^ which have been implicated in ADHD, ASD, MDD, and SCZ^60–63^. *NTM* (**Figure 5C**) regulates the outgrowth of neurites, and is associated with the formation of excitatory synapses^64,65^. It was suggested that haploinsufficiency of *NTM* may influence brain structural volumes and increase the risk for ASD^66,67^. Alterations of *NTM* expression in the dorsolateral prefrontal cortex was also observed in SCZ patients^68^. Of note, *SNX19* at the same locus has been prioritized as a candidate causal gene for SCZ in transcriptomic Mendelian randomization studies^69^. *ANKS1B* (**Figure 5D**) encodes an activity dependent effector protein associated with postsynaptic density^70^, and is involved in long-term depression and synaptic plasticity^71^. *ANKS1B* mutation was found to be enriched in SCZ and ASD^72,73^, and differential methylation was found in *ANKS1B* in prefrontal cortex from SCZ patients^74^. Moreover, *ANKS1B* knockout mice displayed behavior patterns relevant to SCZ including alterations to sensorimotor gating and locomotor activity^71,75^.

Taken together, we have introduced LOGODetect, a scan statistic method to identify local genetic regions showing correlated effects with multiple psychiatric disorders. Complementary to single SNP-based approaches for pleiotropy mapping^17,76^ and genetic correlation estimation methods utilizing genome-wide data^10,20^, our method elucidates the shared genetic architecture between two traits by identifying local genomic segments that are concordant. The candidate genes and regions we identified may be tapping into a set of transdiagnostic mechanisms that underlie all of psychopathology (i.e., the “p” or general factor^39^). In practice, LOGODetect can be used in combination with other methods to further improve statistical power and biological interpretability. For example, it may be of interest to first screen the genome by identifying larger genetic regions^12^ or certain functional annotations^11^ enriched for the shared genetics between two traits. Then, LOGODetect can be applied to these candidate regions to identify the precise genetic segments that explain such sharing. Since high-dimensional sampling remains a challenge, a multi-tier analytical strategy would improve the statistical power and computational burden in the analysis. We believe that LOGODetect has addressed some key limitations in the current practice of cross-trait genetic correlation analysis and will greatly benefit complex trait genetics research.

## Methods

### Genetic Model

Suppose two standardized traits *y*_1_ and *y*_2_ follow the linear model with random effects:

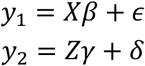

where *X* and *z* are fixed and standardized genotype matrices with *M* columns (i.e. the number of SNPs is *M*); *ϵ* and *δ* are non-genetic effects; *β* and *γ* are *M*-dimensional vectors denoting genetic effects. They follow the multivariate normal distribution:

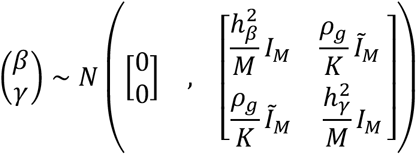

where 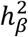 and 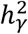 denote the heritability for two traits; *ρ*_*g*_ is the global genetic covariance between two traits; *Ĩ*_*M*_ is a diagonal matrix whose *i*-th diagonal element equals 1 if the effects of the *i*-th SNP on two traits (i.e. *β*_*i*_ and *γ*_*i*_) are correlated and equals 0 if otherwise; *K* is the number of SNPs such that *β*_*i*_ and *γ*_*i*_ are correlated, i.e., *K* = *tr*[*Ĩ*_*M*_]. *β* and *γ* are independent from non-genetic effects *ϵ* and *δ*. The statistical model described here is similar to the polygenic model used in genetic correlation estimation^10^. The difference is that we allow local genetic sharing and do not assume the global genetic covariance to be the same across all the SNPs in the whole genome. Compared to the local genetic correlation estimation method in the literature^12^, we do not assume genetic effects to be fixed. Instead, our framework is a direct generalization of the model developed for global genetic correlation estimation^10,11^. Under the alternative hypothesis, we denote the non-overlapping genetic regions that contribute to multiple traits to be *R*_1_, …, *R*_*r*_ and the union set as 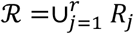 such that *Ĩ*_*M*_[*i*, *i*] = 1 if and only if 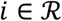. While under the null hypothesis, two traits share no genetic covariance, i.e., 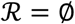.

### Scan Statistic and Scanning Procedure

We use a scan statistics approach to identify regions showing correlated effects between different traits. This type of approach has been used for burden test in a single-trait setting^77^. Suppose *n*_1_, *n*_2_ are the sample sizes for two GWASs, respectively, and we first consider the simpler case that there is no sample overlap between two GWASs. Additionally, we denote the association *z*-scores for two traits as

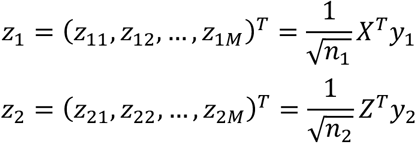

Then, we can define the scan statistic:

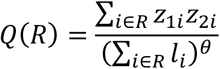

where *R* is the index set for SNPs in a genome region, *l*_*i*_ is the LD score^78^ for the *i*-th SNP, and *θ* is a tuning parameter that controls the strength we penalize over the LD structure. If SNPs in the region *R* show strong, concordant effects on both traits, then the inner product ∑_*i*∈*R*_ *z*_1*i*_*z*_2*i*_ will tend to have a larger absolute value and therefore yield a larger scan statistic. On the contrary, if two traits are genetically independent in the local region, then the corresponding scan statistic would be close to 0. Therefore, the scan statistic is informative to detect local genetic correlation. The purpose of the LD score term in the denominator is to normalize the effect of LD. The expected value of ∑_*i*∈*R*_ *z*_1*i*_*z*_2*i*_ is larger in regions with strong LD. Without the normalization term on the denominator, the method will favor regions with large LD that may not be of biological interest. Further, parameter *θ* affects the size of identified regions. A relatively long segment may not have a large absolute value of scan statistic, due to the penalty in the denominator. A larger *θ* implies stronger penalty, henceforth is more likely to detect smaller signal segments. In particular when *θ* equals 1, |*Q*(*R*)| will attain local maximum with *R* containing only one variant. A reasonable range for *θ* is between 0 and 1, and we used simulations to demonstrate that a *θ* value of 0.5 gives great empirical performance with well-controlled type-I error and reasonable statistical power.

Finally, we use the maximal scan statistic over all possible regions as the test statistic:

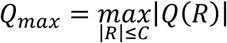

where *C* is a pre-specified parameter that defines the upper boundary of the SNPs count in a region. In practice, *C* can be set based on the number of SNPs in the dataset (e.g. the average number of SNPs in 1 million bases). LOGODetect takes advantages of the flexible framework to scan local regions with varying sizes. Compared to a sliding-window approach based on a pre-specified window size, our method is more appealing since the size of signal region could vary substantially by locus and by trait. We use a Monte Carlo type approach to assess the distribution of *Q*_*max*_ under the null hypothesis. We draw *N* = 5000 pseudo samples 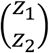 under the null distribution using a procedure detailed in the next section. Then, we estimate the empirical null distribution of *Q*_*max*_ and its 95% upper quantile. Taken together, the scanning procedure works as follows. We scan the genome to find *R*_1_ such that |*Q*(*R*_1_)| reaches the maximum. If |*Q*(*R*_1_)| ≥ *Q*_0.95_, we claim that *R*_1_ is a significant signal region and remove these SNPs from the analysis. Then, we repeat the procedure on the remaining SNPs until no region is declared significant. This procedure controls the family-wise type-I error rate.

### Monte Carlo simulation of pseudo z-score vectors

In order to simulate the null distribution of *Q*_*max*_, we need to generate pseudo z-score vectors. When two GWASs do not have sample overlap, it can be verified that

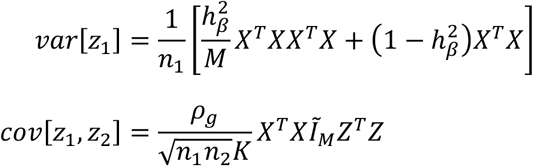

And similarly for *var*[*z*_2_]. Therefore, under *H*_0_, the combined *z*-score vector

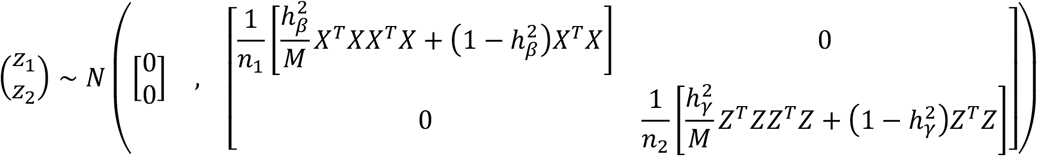

asymptotically. Note that in practice individual genotype data is hard to obtain due to privacy, it is meaningful to analyze based only on summary statistics. Here by using reference panel (e.g. 1000 Genomes Project), 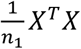 and 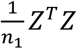 can be estimated as *V*, 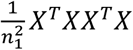 and 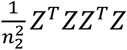 can be estimated as 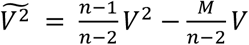, where *n* is the sample size of the reference panel and *V* is the LD matrix of the reference panel. And the genetic heritability for two traits 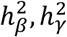 can be estimated through LD score regression^78^. After plugging in the reference LD matrix, we have

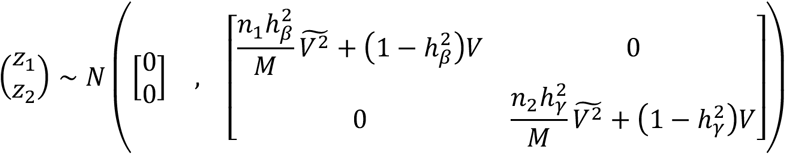

asymptotically under the null.

The random multivariate normal vectors have complex covariance structure, which is computationally challenging as the dimension of the vector can be as high as 10^7^ in GWAS. We developed a computationally tractable method that leverages the LD structure in the genome. First, we split the high-dimensional vector *z* into subvectors *z* = (*z*_(1)_, *z*_(2)_, …, *z*_(*m*)_). Each subvector *z*_(*i*)_ covers SNPs in a 1 MB genome region. We denote the variance matrix of *z* as *Σ* and it can be written as the block matrix form. Denote *Σ*_*i*,*m*_ = *cov*[*z*_(*i*)_, *z*_(*j*)_] as the submatrix of *Σ*, with rows indexed by the *i*-th block *z*_(*i*)_ and columns indexed by the *j*-th block *z*_(*j*)_. Then we use a block-wise tridiagonal matrix to approximate *Σ* by shrinking *Σ*_*i*,*j*_ to 0 if |*i* − *j*| ≥ 2. This approximation is reasonable in the context of GWAS since SNPs should be independent if they are physically apart. Then, we can use an iterative approach to generate each block *z*_(*i*)_ by conditioning on the previous block *z*_(*i*−1)_ via the conditional normal distribution:

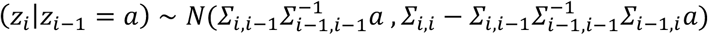

In practice, *Σ*_*i*,*i*_ may be rank deficient and therefore not invertible. We adopt the truncated singular value decomposition (TSVD) method^79^ and use the top *q* singular values and their corresponding singular vectors to calculate the inverse matrix. For numerical stability, we choose *q* to be as large as possible such that the conditional number is less than 1,000^80^. Finally, we standardize each pseudo *z*-score vector so that it has the same mean and variance as the *z*-score vector in real data.

### Extension for sample overlaps

Suppose there are *n*_*s*_ shared samples in the two GWASs, then the linear models can be restated as:

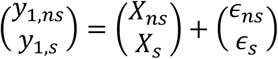

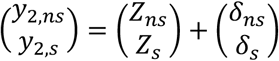

where 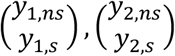 are the standardized phenotypes of all individuals in each GWAS. 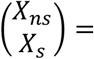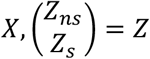 are standardized genotypes of all individuals in each GWAS. *ϵ*_*ns*_, *ϵ*_*s*_, *δ*_*ns*_, *δ*_*s*_ are the non-genetic effects where 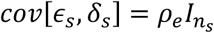. It can be shown that

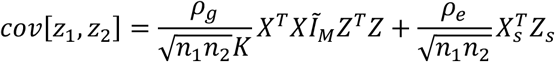

While *var*[*z*_1_] and *var*[*z*_2_] have the same form as no sample overlaps setting. By using reference panel, 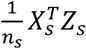 can be replaced by *V*. Therefore, under *H*_0_, the combined *z*-score vectors asymptotically follows multivariate normal distributions

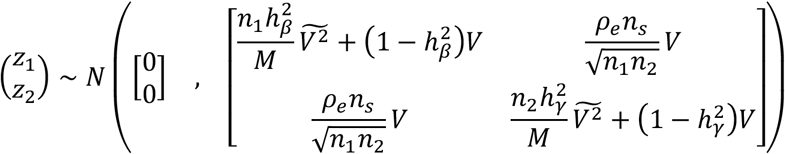

Note that the variance matrix can be split into two terms.

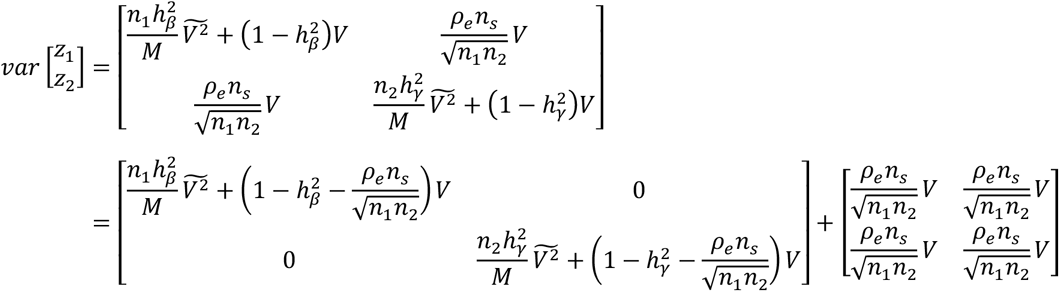

We can independently simulate pseudo samples following the normal distribution with mean 0 and each variance term respectively. Finally, by adding up two vectors simulated with respect to different variance terms, we get the pseudo *z* -score vector of interest. In particular, the parameters 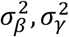, *ρ*_*e*_ ∗ *n*_*s*_ appearing in the *z*-score null distribution are not of our interest, but we need their values while doing Monte Carlo sampling of 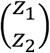. We adopt cross-trait LD score regression^10^ to estimate them. Note that LD score regression is based on random effect random design model setup, which is incompatible with our model assumption, yet we believe it should yield little consequence.

### Genome partition and FDR control

We separated the genome into several LD blocks using ldetect^81^. Each LD block spans 15 MB on average (204 LD blocks in total, **Supplementary Figure 12**). We applied LOGODetect to each LD block separately and identified the local regions with p-value < 0.05 under a family-wise type I error control. We aggregated all the candidate regions across different LD blocks, and applied Benjamini-Hochberg procedure^82^ to control FDR with a cutoff of 0.05.

### Simulation settings

Simulations were based on the genotype data from the WTCCC cohort. We adopted the same quality control procedure as previously described^11^ and only included SNPs on chromosome 1 in the analysis. After quality control, 15,918 individuals and 20,211 SNPs remained in the dataset. Samples were randomly divided into two subsets with equal sample size. We used each subset to simulate the phenotype data.

First, we performed simulations under the null hypothesis to see whether our approach would produce false positive findings. We follow the strict polygenic null, where the effect size level of all the SNPs are the same, and the per-SNP genetic effect was drawn from a normal distribution 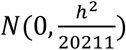 for both traits. To realistically model the polygenic genetic architecture with different levels of genetic effects, we attributed 30% of the trait heritability to 500 randomly chosen SNPs, while the remaining SNPs explain 70% of the total heritability. The per-SNP genetic effect was drawn from a normal distribution 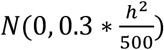 for SNPs with high heritability enrichment, and from 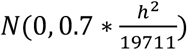 for SNPs with low heritability enrichment. The total heritability *h*^2^ was set to 0.9, 0.3, 0.1, 0.03 and 0.01 for each trait. Each simulation setting was repeated for 1,000 times.

Next, we performed simulations to assess the statistical power. For each trait, we randomly selected *N* = 5 segments, each containing *L* = 100 SNPs, as the signal regions shared between two traits. The genetic effect size for the SNPs in the signal regions follows a multivariate normal distribution

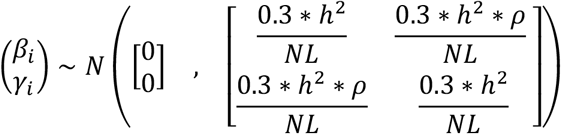

The genetic effect size for the SNPs outside the signal regions follows a different multivariate normal distribution without local genetic correlation

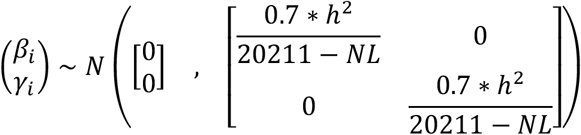

The total heritability *h*^2^ was set to be 0.1 for both traits and the correlation of genetic effect size of two traits *ρ* was set to vary from 0.9 to 0.1. Each simulation setting was repeated 100 times.

### Evaluate model performance

We use three different metrics to quantify the performance of our approach. Denote the true signal segments as *R*_1_, …, *R*_*J*_, and the segments detected by LOGODetect as 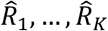. We define the signal points detection rate as the number of true signal SNPs detected by LOGODetect divided by the number of true signal SNPs, that is 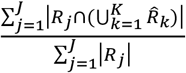. Similarly, we define signal segments detection rate as the number of true signal segments detected by LOGODetect divided by the number of true signal segments, namely 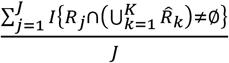, where we call a segment true positive if it overlaps with a true signal segment. Signal points detection rate and signal segments detection rate aim to measure the sensitivity in SNPs level and segments level respectively. To take the extent of the overlap into consideration, we also followed^83^ to define *S*(*R*_*j*_), the G-score with respect to a signal region *R*_*j*_, as 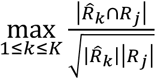 and further define the G-score measure as 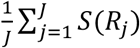. The G-score aims to measure the accuracy and sensitivity together.

### Application of LOGODetect to five psychiatric disorders

We applied LOGODetect to five psychiatric disorders. The European ancestry genotype data from 1000 Genomes Project was used as the reference panel to estimate the LD matrix. For each GWAS data, indels and SNPs not present in the reference panel were removed. The SNPs of minor allele frequency less than 0.01 in the reference panel were also removed. Then for each disorder pair, we filtered out all the strand-ambiguous SNPs and took the overlaps, and we applied LOGODetect to perform the downstream analysis.

### Enrichment analysis

We aggregated 49 non-overlapping segments identified by LOGODetect in five psychiatric disorders and investigated if these segments are enriched in predicted functional regions for a given tissue or cell type. Tissue or cell type-specific functional regions were defined using GenoSkyline-Plus annotations and dichotomized with a cutoff of 0.5. The annotation is robust to the cutoff due to the bimodal pattern in raw GenoSkyline-Plus annotation scores. To assess the statistical significance of enrichment, we randomly selected 49 non-overlapping segments across the genome while matching their sizes with the detected segments, and calculated the overlaps with GenoSkyline-Plus annotations. We repeated the permutation procedure 10,000 times to evaluate the significance of the observed overlap.

We also assessed whether the detected regions were enriched in non-brain tissue types after adjusting for the overlap of brain and non-brain annotations. Specifically, for the pancreatic islets cell type annotation, we removed the annotations that overlap with any of the five significant brain cell type annotations to define the conditional annotation of pancreatic islets. The same procedure was taken to define the conditional annotation of mononuclear cells from peripheral blood. Afterwards, permutation test was performed on these two conditional annotations.

### URLs

Summary statistics data of five psychiatric disorder can be downloaded on the PGC website, http://www.med.unc.edu/pgc/downloads; 66 GenoSkyline-Plus cell-type specific functional annotations, http://genocanyon.med.yale.edu/GenoSkyline; Fetal brain TAD data https://www.ncbi.nlm.nih.gov/geo/query/acc.cgi?acc=GSE77565; Adult brain TAD data http://resource.psychencode.org.

### Code availability

LOGODetect software is available at https://github.com/ghm17/LOGODetect.

## Acknowledgements

We thank Dr. Daifeng Wang for sharing the adult brain TAD data generated by the PsychENCODE consortium. This study makes use of summary statistics from the Psychiatric Genomics Consortium. We thank the investigators for providing publicly accessible GWAS summary statistics. Li and Lu acknowledge research support from the Waisman Center pilot grant program at University of Wisconsin-Madison. Hou acknowledges research support from the National Science Foundation of China (Grant No. 11601259) and Shanghai Municipal Science and Technology Major Project (Grant No. 2017SHZDZX01).

## Author contribution

H.G., Q.L. and L.H. designed the idea. H.G. performed data analysis. H.G. developed the software. H.G., J.J.L, Q.L. and L.H. interpreted the results. H.G., Q.L. and L.H. drafted the manuscript.

## Competing financial interests

The authors declare no competing financial interests.

## Notes

https://github.com/ghm17/LOGODetect

http://www.med.unc.edu/pgc/downloads

http://genocanyon.med.yale.edu/GenoSkyline

https://www.ncbi.nlm.nih.gov/geo/query/acc.cgi?acc=GSE77565

http://resource.psychencode.org

## References

1. Sivakumaran, S. et al. Abundant pleiotropy in human complex diseases and traits. Am J Hum Genet 89, 607–18 (2011).

2. Solovieff, N., Cotsapas, C., Lee, P.H., Purcell, S.M. & Smoller, J.W. Pleiotropy in complex traits: challenges and strategies. Nat Rev Genet 14, 483–95 (2013).

3. van Rheenen, W., Peyrot, W.J., Schork, A.J., Lee, S.H. & Wray, N.R. Genetic correlations of polygenic disease traits: from theory to practice. Nature Reviews Genetics, 1 (2019).

4. Andreassen, O.A. et al. Improved detection of common variants associated with schizophrenia by leveraging pleiotropy with cardiovascular-disease risk factors. Am J Hum Genet 92, 197–209 (2013).

5. Yang, W.H. et al. Recurrent infection progressively disables host protection against intestinal inflammation. Science 358(2017).

6. Liang, J. et al. Single-trait and multi-trait genome-wide association analyses identify novel loci for blood pressure in African-ancestry populations. PLoS genetics 13, e1006728 (2017).

7. Andreassen, O.A. et al. Genetic pleiotropy between multiple sclerosis and schizophrenia but not bipolar disorder: differential involvement of immune-related gene loci. Molecular psychiatry 20, 207 (2015).

8. Krapohl, E. et al. Multi-polygenic score approach to trait prediction. Molecular psychiatry 23, 1368 (2018).

9. Maier, R. et al. Joint analysis of psychiatric disorders increases accuracy of risk prediction for schizophrenia, bipolar disorder, and major depressive disorder. The American Journal of Human Genetics 96, 283–294 (2015).

10. Bulik-Sullivan, B. et al. An atlas of genetic correlations across human diseases and traits. Nature genetics 47, 1236 (2015).

11. Lu, Q. et al. A Powerful Approach to Estimating Annotation-Stratified Genetic Covariance via GWAS Summary Statistics. Am J Hum Genet 101, 939–964 (2017).

12. Shi, H., Mancuso, N., Spendlove, S. & Pasaniuc, B. Local genetic correlation gives insights into the shared genetic architecture of complex traits. The American Journal of Human Genetics 101, 737–751 (2017).

13. Brainstorm, C. et al. Analysis of shared heritability in common disorders of the brain. Science 360(2018).

14. Turley, P. et al. Multi-trait analysis of genome-wide association summary statistics using MTAG. Nature genetics 50, 229 (2018).

15. Majumdar, A., Haldar, T., Bhattacharya, S. & Witte, J.S. An efficient Bayesian meta-analysis approach for studying cross-phenotype genetic associations. PLoS genetics 14, e1007139 (2018).

16. Zhou, X. & Stephens, M. Genome-wide efficient mixed-model analysis for association studies. Nature genetics 44, 821 (2012).

17. Pickrell, J.K. et al. Detection and interpretation of shared genetic influences on 42 human traits. Nature genetics 48, 709 (2016).

18. Yang, J. et al. Common SNPs explain a large proportion of the heritability for human height. Nat Genet 42, 565–9 (2010).

19. Boyle, E.A., Li, Y.I. & Pritchard, J.K. An Expanded View of Complex Traits: From Polygenic to Omnigenic. Cell 169, 1177–1186 (2017).

20. Lee, S.H., Yang, J., Goddard, M.E., Visscher, P.M. & Wray, N.R. Estimation of pleiotropy between complex diseases using single-nucleotide polymorphism-derived genomic relationships and restricted maximum likelihood. Bioinformatics 28, 2540–2542 (2012).

21. Vattikuti, S., Guo, J. & Chow, C.C. Heritability and genetic correlations explained by common SNPs for metabolic syndrome traits. PLoS genetics 8, e1002637 (2012).

22. Pasaniuc, B. & Price, A.L. Dissecting the genetics of complex traits using summary association statistics. Nature Reviews Genetics 18, 117 (2017).

23. Kessler, R.C., Chiu, W.T., Demler, O. & Walters, E.E. Prevalence, Severity, and Comorbidity of 12-Month DSM-IV Disorders in the National Comorbidity Survey Replication. JAMA Psychiatry 62, 617–627 (2005).

24. Larsson, H. et al. Risk of bipolar disorder and schizophrenia in relatives of people with attention-deficit hyperactivity disorder. The British Journal of Psychiatry 203, 103–106 (2013).

25. Consortium, C.-D.G.o.t.P.G. Identification of risk loci with shared effects on five major psychiatric disorders: a genome-wide analysis. The Lancet 381, 1371–1379 (2013).

26. Lee, S.H. et al. Genetic relationship between five psychiatric disorders estimated from genome-wide SNPs. Nature genetics 45, 984 (2013).

27. Gratten, J., Wray, N.R., Keller, M.C. & Visscher, P.M. Large-scale genomics unveils the genetic architecture of psychiatric disorders. Nature neuroscience 17, 782 (2014).

28. Smoller, J.W. & Finn, C.T. Family, twin, and adoption studies of bipolar disorder. in American Journal of Medical Genetics Part C: Seminars in Medical Genetics Vol. 123 48–58 (Wiley Online Library, 2003).

29. Lichtenstein, P., Carlström, E., Råstam, M., Gillberg, C. & Anckarsäter, H. The genetics of autism spectrum disorders and related neuropsychiatric disorders in childhood. American Journal of Psychiatry 167, 1357–1363 (2010).

30. Cole, J., Ball, H.A., Martin, N.C., Scourfield, J. & Mcguffin, P. Genetic overlap between measures of hyperactivity/inattention and mood in children and adolescents. Journal of the American Academy of Child & Adolescent Psychiatry 48, 1094–1101 (2009).

31. Rommelse, N.N., Franke, B., Geurts, H.M., Hartman, C.A. & Buitelaar, J.K. Shared heritability of attention-deficit/hyperactivity disorder and autism spectrum disorder. European child & adolescent psychiatry 19, 281–295 (2010).

32. Stahl, E.A. et al. Genome-wide association study identifies 30 loci associated with bipolar disorder. Nature genetics 51, 793 (2019).

33. Pardiñas, A.F. et al. Common schizophrenia alleles are enriched in mutation-intolerant genes and in regions under strong background selection. Nature genetics 50, 381 (2018).

34. Wray, N.R. et al. Genome-wide association analyses identify 44 risk variants and refine the genetic architecture of major depression. Nature genetics 50, 668 (2018).

35. Demontis, D. et al. Discovery of the first genome-wide significant risk loci for attention deficit/hyperactivity disorder. Nature genetics 51, 63 (2019).

36. Grove, J. et al. Identification of common genetic risk variants for autism spectrum disorder. Nature genetics 51, 431 (2019).

37. Lichtenstein, P. et al. Common genetic determinants of schizophrenia and bipolar disorder in Swedish families: a population-based study. The Lancet 373, 234–239 (2009).

38. Pettersson, E., Larsson, H. & Lichtenstein, P. Common psychiatric disorders share the same genetic origin: a multivariate sibling study of the Swedish population. Molecular psychiatry 21, 717 (2016).

39. Grotzinger, A.D. et al. Genomic structural equation modelling provides insights into the multivariate genetic architecture of complex traits. Nature human behaviour 3, 513 (2019).

40. Ghaziuddin, M., Ghaziuddin, N. & Greden, J. Depression in persons with autism: Implications for research and clinical care. Journal of autism and developmental disorders 32, 299–306 (2002).

41. Lu, Q. et al. Systematic tissue-specific functional annotation of the human genome highlights immune-related DNA elements for late-onset Alzheimer’s disease. PLoS genetics 13, e1006933 (2017).

42. Won, H. et al. Chromosome conformation elucidates regulatory relationships in developing human brain. Nature 538, 523 (2016).

43. Wang, D. et al. Comprehensive functional genomic resource and integrative model for the human brain. Science 362, eaat8464 (2018).

44. Hyde, C.L. et al. Identification of 15 genetic loci associated with risk of major depression in individuals of European descent. Nature genetics 48, 1031 (2016).

45. Lionel, A.C. et al. Rare copy number variation discovery and cross-disorder comparisons identify risk genes for ADHD. Science translational medicine 3, 95ra75–95ra75 (2011).

46. McClay, J.L. et al. Genome-wide pharmacogenomic analysis of response to treatment with antipsychotics. Molecular psychiatry 16, 76 (2011).

47. McClay, J.L. et al. Genome-wide pharmacogenomic study of neurocognition as an indicator of antipsychotic treatment response in schizophrenia. Neuropsychopharmacology 36, 616 (2011).

48. Flanagan, J.G. & Vanderhaeghen, P. The ephrins and Eph receptors in neural development. Annual review of neuroscience 21, 309–345 (1998).

49. Himanen, J.-P. et al. Repelling class discrimination: ephrin-A5 binds to and activates EphB2 receptor signaling. Nature neuroscience 7, 501 (2004).

50. Murai, K.K. & Pasquale, E.B. Eph receptors and ephrins in neuron–astrocyte communication at synapses. Glia 59, 1567–1578 (2011).

51. Cooper, M.A., Kobayashi, K. & Zhou, R. Ephrin-A5 regulates the formation of the ascending midbrain dopaminergic pathways. Developmental neurobiology 69, 36–46 (2009).

52. Robinson, D.L., Zitzman, D.L. & Williams, S.K. Mesolimbic dopamine transients in motivated behaviors: focus on maternal behavior. Frontiers in psychiatry 2, 23 (2011).

53. Sheleg, M., Yochum, C.L., Wagner, G.C., Zhou, R. & Richardson, J.R. Ephrin-A5 deficiency alters sensorimotor and monoaminergic development. Behavioural brain research 236, 139–147 (2013).

54. Maletic, V. et al. Neurobiology of depression: an integrated view of key findings. International journal of clinical practice 61, 2030–2040 (2007).

55. Parkitna, J.R. et al. Loss of the serum response factor in the dopamine system leads to hyperactivity. The FASEB Journal 24, 2427–2435 (2010).

56. Breiderhoff, T. et al. Sortilin-related receptor SORCS3 is a postsynaptic modulator of synaptic depression and fear extinction. PLoS One 8, e75006 (2013).

57. Dark, C., Homman-Ludiye, J. & Bryson-Richardson, R.J. The role of ADHD associated genes in neurodevelopment. Developmental biology 438, 69–83 (2018).

58. Christiansen, G.B. et al. The sorting receptor SorCS3 is a stronger regulator of glutamate receptor functions compared to GABAergic mechanisms in the hippocampus. Hippocampus 27, 235–248 (2017).

59. Oetjen, S., Mahlke, C., Hermans-Borgmeyer, I. & Hermey, G. Spatiotemporal expression analysis of the growth factor receptor SorCS3. Journal of Comparative Neurology 522, 3386–3402 (2014).

60. Dev, K.K. & Henley, J.M. The schizophrenic faces of PICK1. Trends in pharmacological sciences 27, 574–579 (2006).

61. Guney, E. et al. Serum nerve growth factor (NGF) levels in children with attention deficit/hyperactivity disorder (ADHD). Neuroscience letters 560, 107–111 (2014).

62. Kajizuka, M. et al. Serum levels of platelet-derived growth factor BB homodimers are increased in male children with autism. Progress in Neuro-Psychopharmacology and Biological Psychiatry 34, 154–158 (2010).

63. Wiener, C.D. et al. Serum levels of nerve growth factor (NGF) in patients with major depression disorder and suicide risk. Journal of affective disorders 184, 245–248 (2015).

64. Gil, O.D., Zanazzi, G., Struyk, A.F. & Salzer, J.L. Neurotrimin mediates bifunctional effects on neurite outgrowth via homophilic and heterophilic interactions. Journal of Neuroscience 18, 9312–9325 (1998).

65. Chen, S. et al. Neurotrimin expression during cerebellar development suggests roles in axon fasciculation and synaptogenesis. Journal of neurocytology 30, 927–937 (2001).

66. Maruani, A. et al. 11q24. 2-25 micro-rearrangements in autism spectrum disorders: Relation to brain structures. American Journal of Medical Genetics Part A 167, 3019–3030 (2015).

67. Minhas, H.M. et al. An unbalanced translocation involving loss of 10q26. 2 and gain of 11q25 in a pedigree with autism spectrum disorder and cerebellar juvenile pilocytic astrocytoma. American Journal of Medical Genetics Part A 161, 787–791 (2013).

68. Karis, K. et al. Altered expression profile of IgLON family of neural cell adhesion molecules in the dorsolateral prefrontal cortex of schizophrenic patients. Frontiers in molecular neuroscience 11, 8 (2018).

69. Zhu, Z. et al. Integration of summary data from GWAS and eQTL studies predicts complex trait gene targets. Nature genetics 48, 481 (2016).

70. Ghersi, E., Vito, P., Lopez, P., Abdallah, M. & D’Adamio, L. The intracellular localization of amyloid β protein precursor (AβPP) intracellular domain associated protein-1 (AIDA-1) is regulated by AβPP and alternative splicing. Journal of Alzheimer's Disease 6, 67–78 (2004).

71. Tindi, J.O. et al. ANKS1B gene product AIDA-1 controls hippocampal synaptic transmission by regulating GluN2B subunit localization. Journal of Neuroscience 35, 8986–8996 (2015).

72. Purcell, S.M. et al. A polygenic burden of rare disruptive mutations in schizophrenia. Nature 506, 185 (2014).

73. Li, J. et al. Integrated systems analysis reveals a molecular network underlying autism spectrum disorders. Molecular systems biology 10(2014).

74. Han, S. et al. Integrating brain methylome with GWAS for psychiatric risk gene discovery. bioRxiv, 440206 (2018).

75. Enga, R.M. et al. Initial characterization of behavior and ketamine response in a mouse knockout of the post-synaptic effector gene Anks1b. Neuroscience letters 641, 26–32 (2017).

76. Giambartolomei, C. et al. Bayesian test for colocalisation between pairs of genetic association studies using summary statistics. PLoS genetics 10, e1004383 (2014).

77. Li, Z. et al. Dynamic Scan Procedure for Detecting Rare-Variant Association Regions in Whole-Genome Sequencing Studies. The American Journal of Human Genetics 104, 802–814 (2019).

78. Bulik-Sullivan, B.K. et al. LD Score regression distinguishes confounding from polygenicity in genome-wide association studies. Nature genetics 47, 291 (2015).

79. Hansen, P.C. Truncated singular value decomposition solutions to discrete ill-posed problems with ill-determined numerical rank. SIAM Journal on Scientific and Statistical Computing 11, 503–518 (1990).

80. Won, J.H., Lim, J., Kim, S.J. & Rajaratnam, B. Condition - number - regularized covariance estimation. Journal of the Royal Statistical Society: Series B (Statistical Methodology) 75, 427–450 (2013).

81. Berisa, T. & Pickrell, J.K. Approximately independent linkage disequilibrium blocks in human populations. Bioinformatics 32, 283 (2016).

82. Benjamini, Y. & Hochberg, Y. Controlling the false discovery rate: a practical and powerful approach to multiple testing. Journal of the Royal statistical society: series B (Methodological) 57, 289–300 (1995).

83. Jeng, X.J., Cai, T.T. & Li, H. Simultaneous discovery of rare and common segment variants. Biometrika 100, 157–172 (2012).

