## Supplementary Figure for "Detecting Local Genetic Correlations with Scan Statistics"

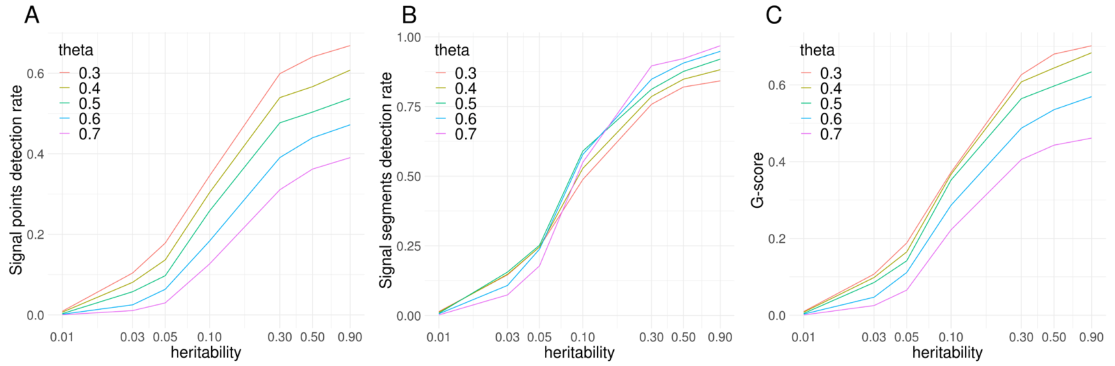


Fig S1. Power evaluation of LOGODetect. (A-C) show statistical power assessed by three measures: Signal points detection rate, Signal segments detection rate and G-score. Horizontal axis is in log scale. Heritability represents the total heritability for both traits. For each trait, we randomly choose N=5 segments, each contains L=100 SNPs, as the signal regions. The heritability for the high enrichment SNPs set is set to be 30% total heritability. The correlation of genetic effect size of two traits ρ is set to be 0.9. Each simulation setting is repeated for 100 times.

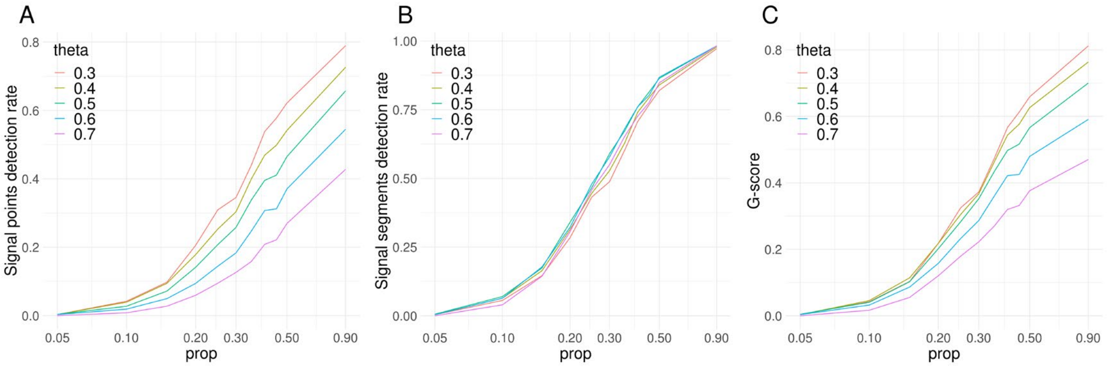


Fig S2. Power evaluation of LOGODetect. (A-C) show statistical power assessed by three measures: Signal points detection rate, Signal segments detection rate and G-score. Horizontal axis is in log scale. Prop represents the proportion of the high enrichment SNP sets heritability divided by the total heritability. For each trait, we randomly choose N=5 segments, each contains L=100 SNPs, as the signal regions. The total heritability is set to be 0.1 for both traits. The correlation of genetic effect size of two traits ρ is set to be 0.9. Each simulation setting is repeated for 100 times.

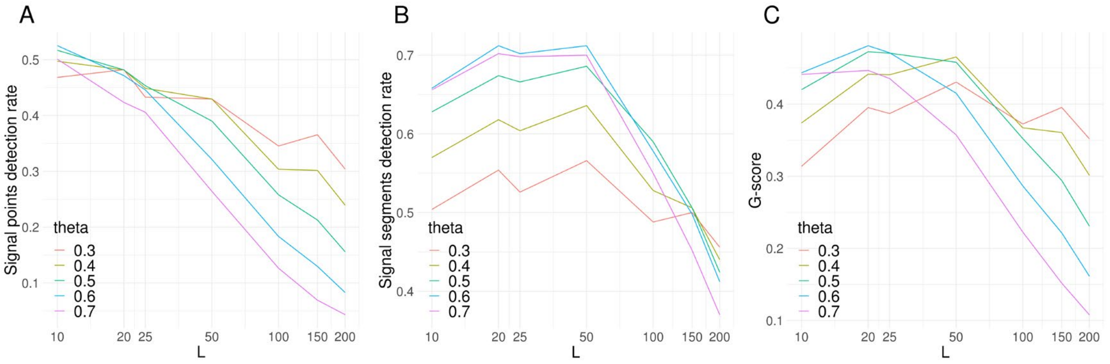


Fig S3. Power evaluation of LOGODetect. (A-C) show statistical power assessed by three measures: Signal points detection rate, Signal segments detection rate and G-score. Horizontal axis is in log scale. L represents the length of true signal region. For each trait, we randomly choose N=5 segments, each contains L SNPs, as the signal regions. The total heritability is set to be 0.1 for both traits. The heritability for the high enrichment SNPs set is set to be 30% total heritability. The correlation of genetic effect size of two traits ρ is set to be 0.9. Each simulation setting is repeated for 100 times.

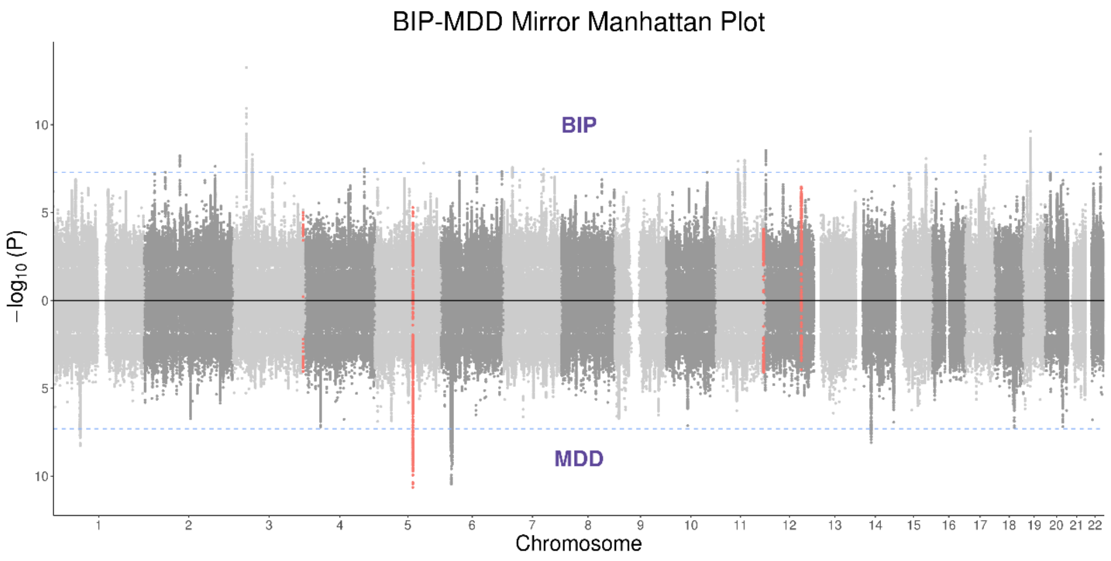


Fig S4. Mirror Manhattan plot for BIP-MDD, red dots represent SNPs located in the LOGODetect detected regions.

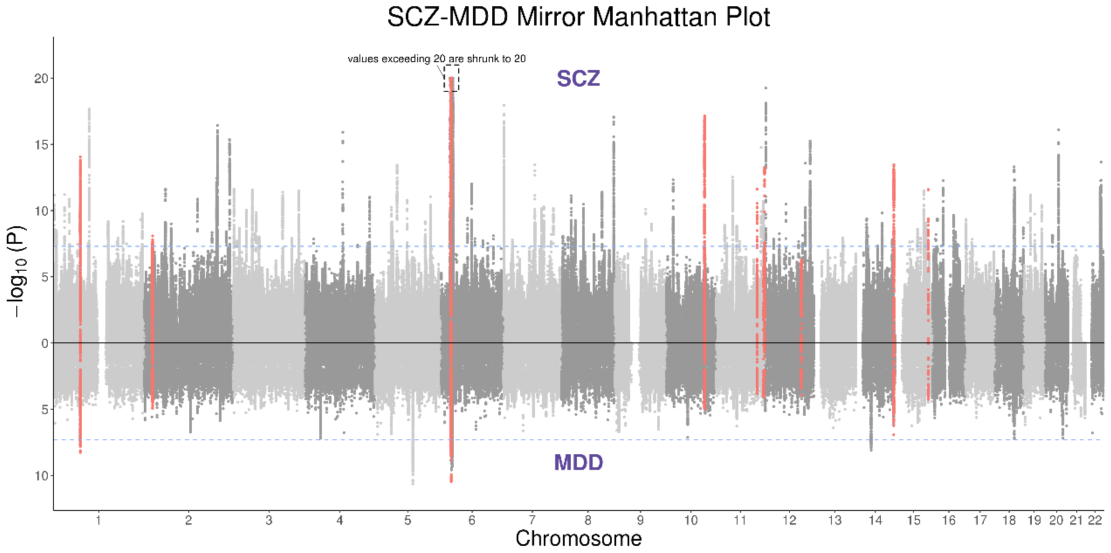


Fig S5. Mirror Manhattan plot for SCZ-MDD, red dots represent SNPs located in the LOGODetect detected regions. For SCZ, one locus on chromosome 6 have $-\log_{10} P$ value exceeding 20, those values are shrunk to 20 for conciseness.

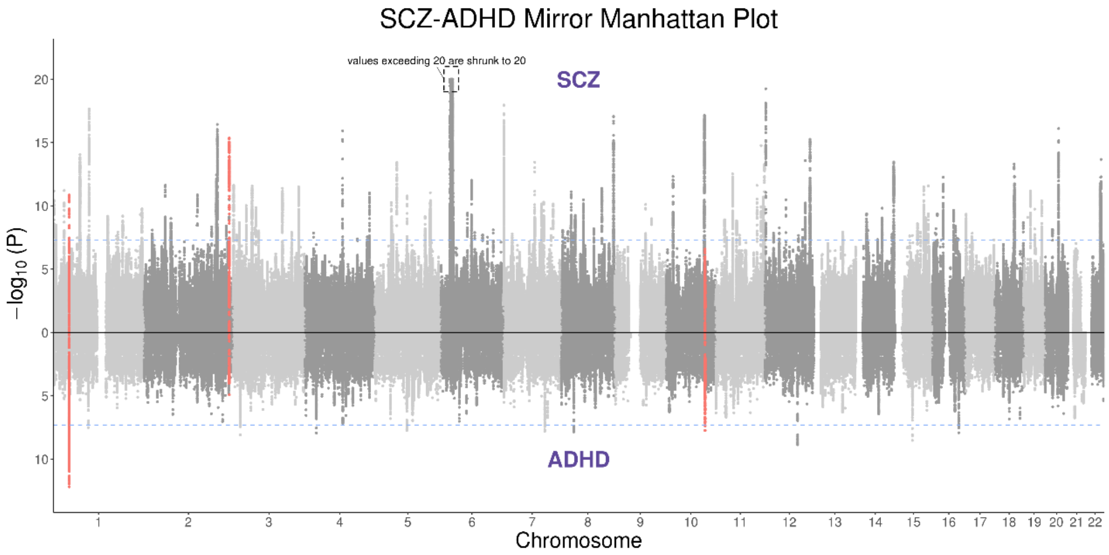


Fig S6. Mirror Manhattan plot for SCZ-ADHD, red dots represent SNPs located in the LOGODetect detected regions. For SCZ, one locus on chromosome 6 have $-\log_{10} P$ value exceeding 20, those values are shrunk to 20 for conciseness.

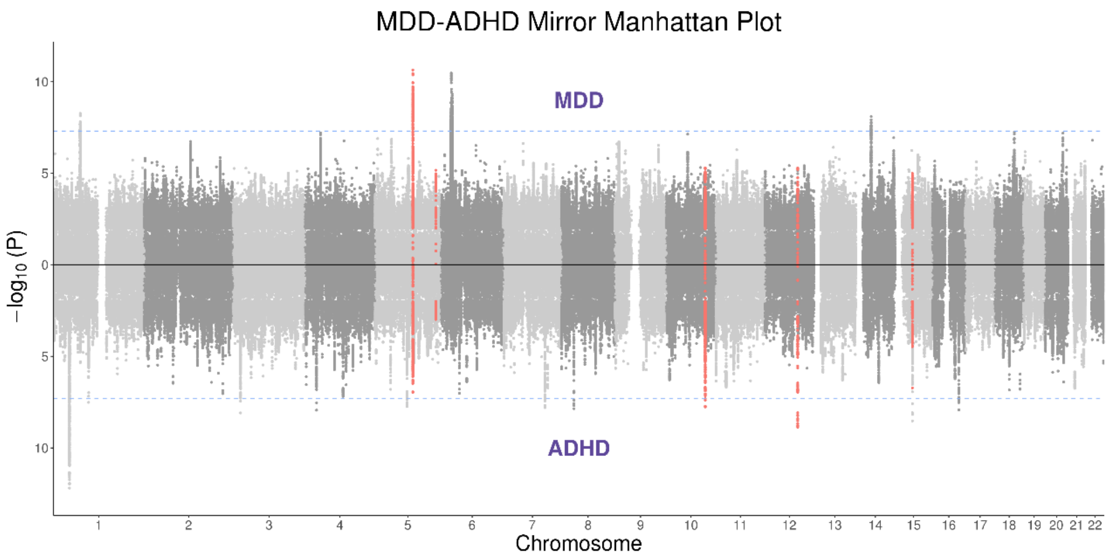


Fig S7. Mirror Manhattan plot for MDD-ADHD, red dots represent SNPs located in the LOGODetect detected regions.

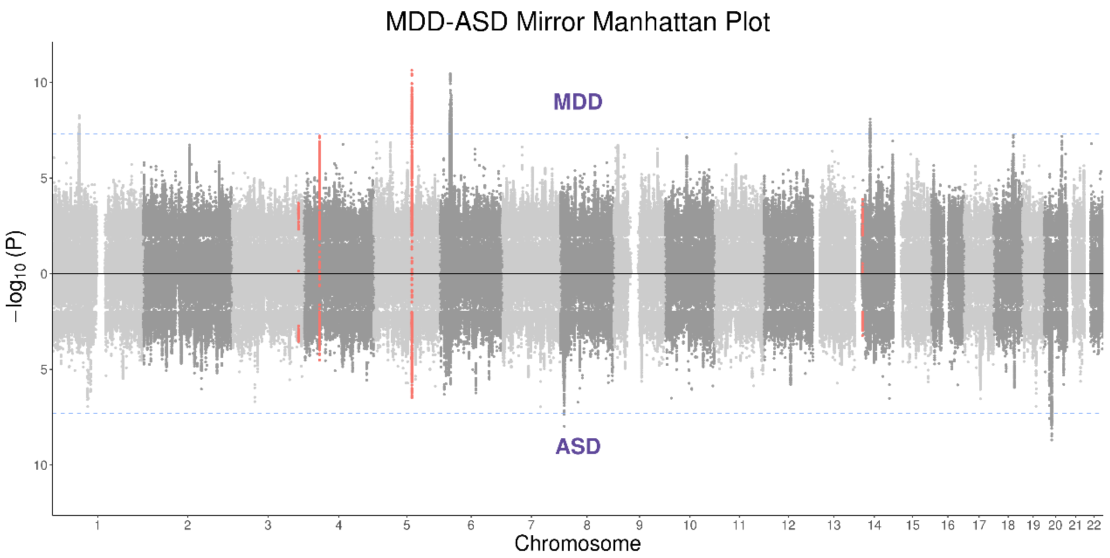


Fig S8. Mirror Manhattan plot for MDD-ASD, red dots represent SNPs located in the LOGODetect detected regions.

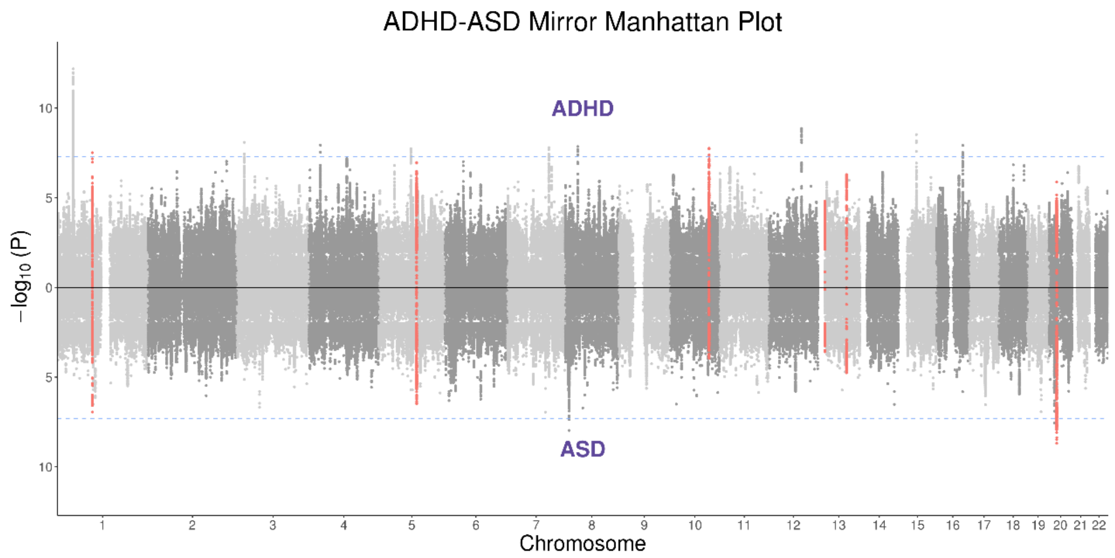


Fig S9. Mirror Manhattan plot for ADHD-ASD, red dots represent SNPs located in the LOGODetect detected regions.

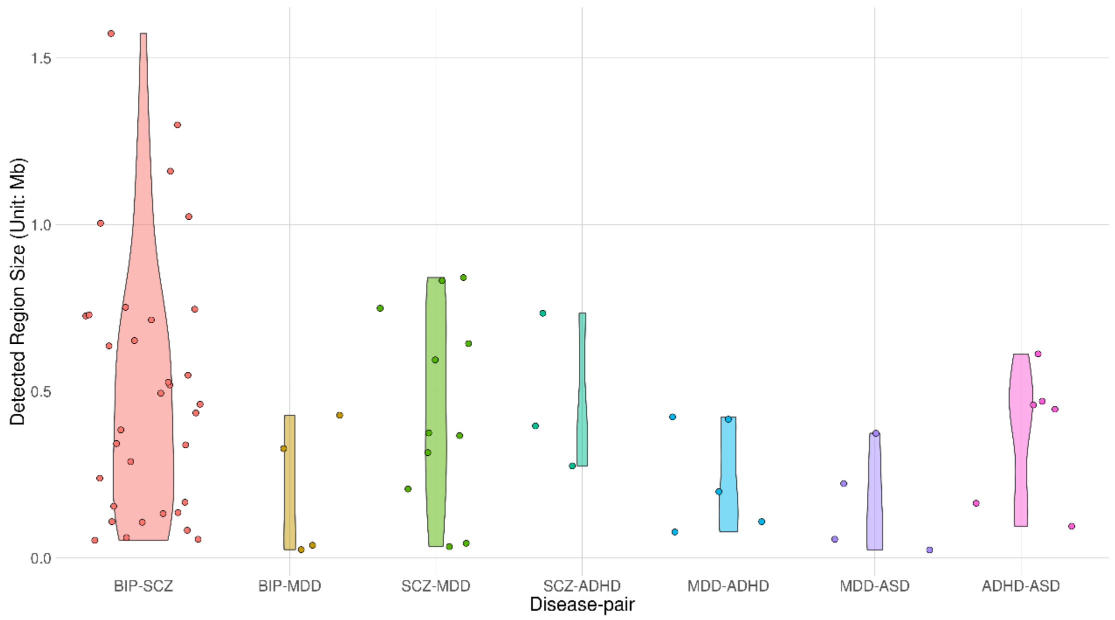


Fig S10. Violin plot of detected region size in different disease-pairs. Only 7 disease-pairs having detected significant regions are shown. Each violin area is scaled proportionally to the number of detected segments.

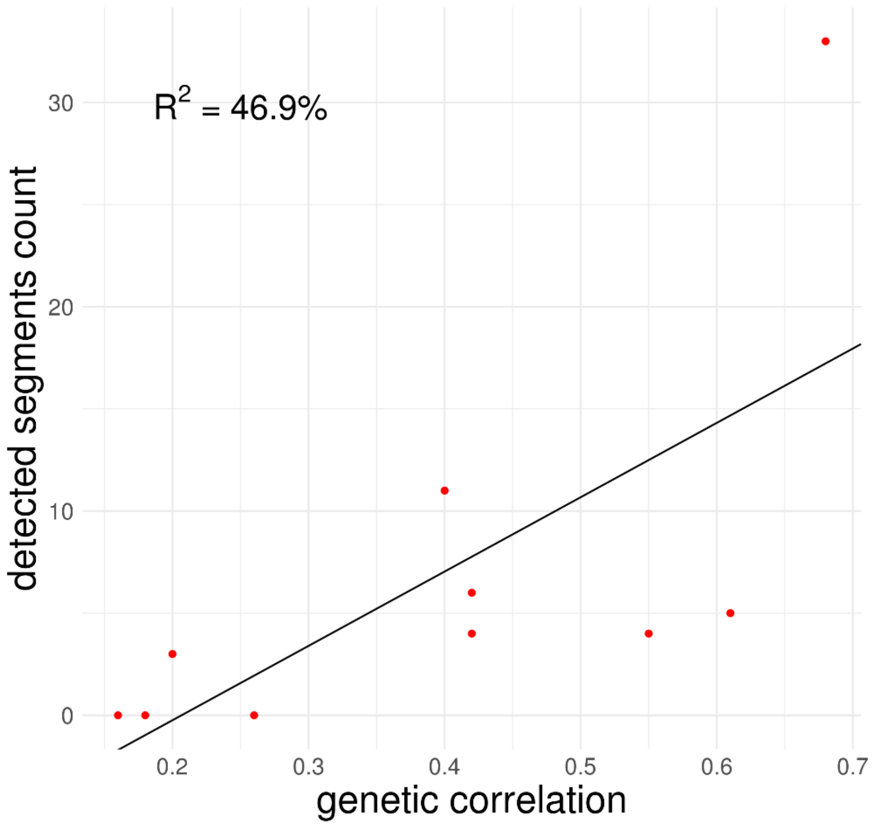


Figure S11: LOGODetect detected segments count and genetic correlation estimated by cross-trait LDSC are concordant, the larger the genetic correlation, the more segments detected by LOGODetect. The R-suqared equals 46.9%.

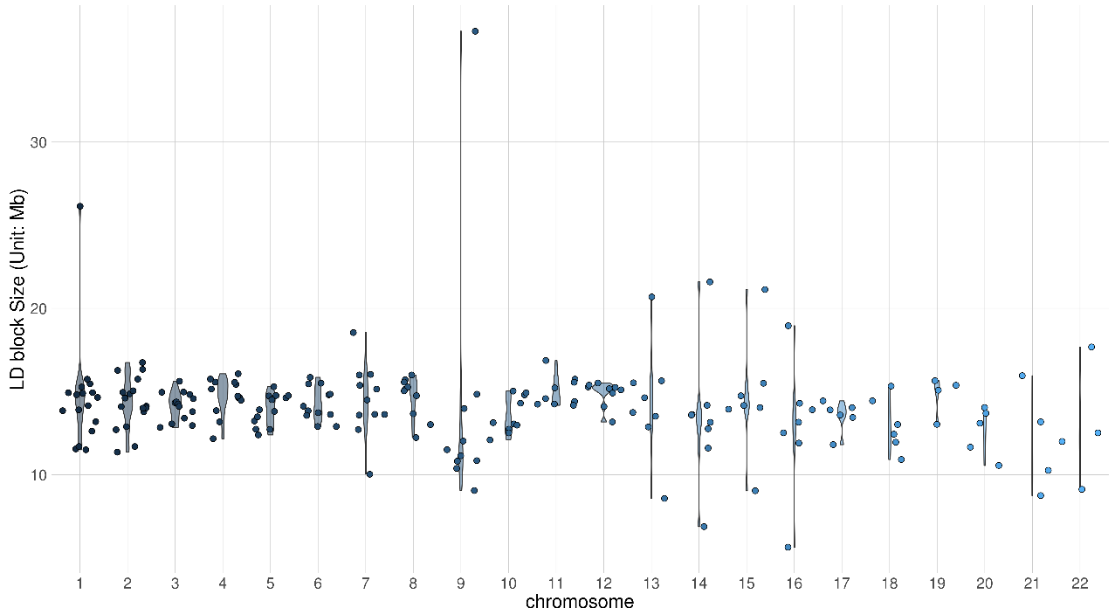


Fig S12. 204 LD blocks size distribution.
